# *LMNA* Haploinsufficiency in Human iPSC-Derived Cardiac Organoids Reveals Early Fibrotic Signaling as a Therapeutically Targetable Process

**DOI:** 10.64898/2026.03.25.714182

**Authors:** A. Zuniga, R. Dulce, K. D. Asensi, A. Chakraborty, B. DeRosa, P. Levitan, S. Borges, R. Volonterio, M. Lopez, J. Dollar, G. Yenisehirli, S. Rodriguez, B. Schachner, D. Dykxhoorn, J. Hare, S. Kurtenbach

## Abstract

*LMNA* mutations are a major cause of dilated cardiomyopathy (DCM), with haploinsufficiency representing a common pathogenic mechanism. Yet the earliest disease-initiating events remain poorly defined. Here, we identify a novel intronic splice-site variant, c.937-1G>A, that disrupts pre-mRNA splicing, induces nonsense-mediated mRNA decay, and results in *LMNA* haploinsufficiency. We differentiated induced pluripotent stem cells (iPSCs) generated from a patient’s peripheral blood mononuclear cells (PBMCs) carrying this mutation into self-patterning human cardiac organoids to define the early consequences of reduced *LMNA* dosage in a multicellular human context. Single-nucleus transcriptomics revealed broad multicellular remodeling beyond the previously implicated fibroblast and cardiomyocyte populations. Functionally, *LMNA*-mutant organoids exhibited altered calcium handling and impaired contractility. These changes were accompanied by increased reactive oxygen species (ROS) and early pro-fibrotic signaling characterized by increased periostin (POSTN) secretion, CTGF expression, and increased p38 MAPK phosphorylation. Treatment with the antifibrotic drug nintedanib attenuated this response, partially rescued rhythmic beating, reduced POSTN secretion and CTGF expression, and broadly suppressed fibrosis-associated transcriptional programs. Together, these findings show that *LMNA* haploinsufficiency initiates multicellular cardiac remodeling and early targetable profibrotic signaling during human cardiac development, providing mechanistic insight into the earliest stages of *LMNA*-associated cardiomyopathy and establishing a human cardiac organoid platform for therapeutic discovery.

## INTRODUCTION

The most prevalent cardiac phenotype associated with *LMNA* mutations is dilated cardiomyopathy (DCM), a major cause of heart failure characterized by ventricular chamber dilation and impaired systolic function [1]. *LMNA*-associated DCM (*LMNA*-DCM) accounts for approximately 4-8% of familial DCM cases and represents one of the most malignant inherited cardiomyopathies, exhibiting high penetrance, autosomal dominant inheritance, and rapid disease progression with early mortality [2–4]. The *LMNA* gene encodes the intermediate filament proteins Lamin A and Lamin C, which are key structural components of the nuclear lamina, a protein meshwork lining the inner nuclear membrane that regulates nuclear architecture, chromatin organization, and gene expression [5]. Mutations in *LMNA*, collectively termed laminopathies, display substantial allelic heterogeneity, with mutation location and type influencing disease phenotype and severity.

Much of our current understanding of *LMNA*-associated cardiomyopathy is derived from transgenic mouse models, most notably the *LMNA*^H222P/H222P^ model, which recapitulates several hallmark features of the disease, including chamber dilation, conduction abnormalities, myocardial fibrosis, and activation of stress-responsive signaling pathways [6]. However, these models do not fully reflect the heterozygous genetic context observed in *LMNA*-DCM patients. Moreover, interspecies differences in nuclear lamina biology, cardiac development, and fibrotic remodeling limit murine systems from recapitulating early human-specific disease mechanisms.

Induced pluripotent stem cell (iPSC)-derived cardiomyocytes have emerged as an important platform for modeling *LMNA* mutations and have provided insights into cellular phenotypes, such as nuclear abnormalities, contractile dysfunction, and impaired calcium handling [7, 8]. However, these systems lack the multicellular complexity required to elucidate tissue-level interactions and remodeling processes that shape disease progression. This limitation is especially relevant to fibrosis, a prominent feature of *LMNA*-DCM that arises through coordinated signaling across multiple cardiac cell types. Consequently, the cellular events by which reduced *LMNA* dosage initiates cardiac remodeling, particularly before the development of overt structural disease, remain poorly understood.

In this study, we used three-dimensional, self-patterning cardiac organoids derived from patient-specific iPSCs and an isogenic gene-corrected control to investigate the earliest pathogenic consequences of *LMNA* haploinsufficiency caused by a novel intronic *LMNA* splice-site mutation (c.937-1G>A) that induces nonsense-mediated mRNA decay. We demonstrate that reduced *LMNA* dosage is sufficient to initiate broad, lineage-specific remodeling across multiple cardiac cell populations, with prominent functional and electrophysiological abnormalities accompanied by activation of early profibrotic transcriptional programs. Furthermore, we identify early profibrotic signaling as a pharmacologically targetable aspect of this phenotype, establishing cardiac organoids as a platform for investigating early disease mechanisms and therapeutic responses in *LMNA*-DCM.

## RESULTS

### A Novel Intronic *LMNA* c.937-1G>A Mutation Causes *LMNA* Haploinsufficiency via Nonsense-Mediated Decay

We previously identified a heterozygous *LMNA* mutation (c.937-1G>A) in a patient enrolled in the POSEIDON-DCM clinical trial who was diagnosed with heart failure secondary to dilated cardiomyopathy and had received a biventricular pacemaker [9]. The patient presented with clinical features consistent with DCM, including reduced left ventricular ejection fraction (LVEF <35%), atrial arrhythmias, and mitral valve regurgitation. Review of the family history revealed a pattern of inherited cardiac disease, with the patient’s mother succumbing to heart failure, a maternal uncle suffering from myocardial infarction, and two maternal cousins diagnosed with heart failure (Figure 1A). Bioinformatic analysis indicated that the c.937-1G>A variant disrupts the 3′ splice acceptor site of intron 5 of the *LMNA* gene. To model the consequences of this variant, peripheral blood mononuclear cells (PBMCs) from the patient were reprogrammed into induced pluripotent stem cells (iPSCs) using a non-integrating Sendai virus system. Sanger sequencing confirmed preservation of the heterozygous (c.937-1G>A) mutation following reprogramming (Figure 1B). Characterization and quality control of the iPSCs confirmed absence of residual Sendai virus, a normal karyotype, and pluripotency through expression of SOX2, SSEA4, and OCT4, as well as trilineage differentiation potential (Supplementary Figure 1A-E). Subsequently, we generated a CRISPR-corrected iPSC line to enable isogenic comparisons (Figure 1B). These iPSCs exhibited normal morphology and growth (Supplementary Figure 1C), and both lines efficiently differentiated into beating cardiac organoids (Supplementary Video). CRISPR correction of the *LMNA* variant led to a marked increase in *LMNA* RNA and protein levels in cardiac organoids (Figure 1C-E), consistent with reduced expression in the mutant line.

**Figure 1.**
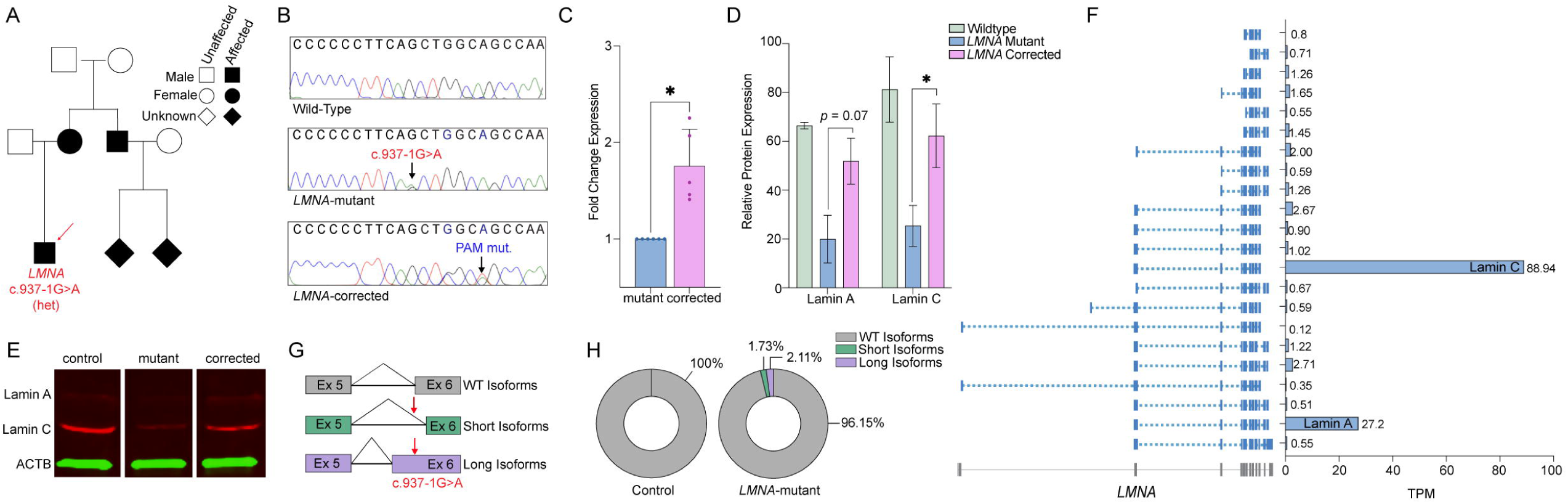
*LMNA* Splice-Site Mutation Causes Haploinsufficiency and is Rescued by Genetic Correction. **(A)** Pedigree of the affected family. The individual sequenced and from which the PBMCs were used to generate the iPSCs is highlighted (red arrow). **(B)** Sanger sequencing confirming retention of the heterozygous *LMNA* c.937-1G mutation in patient cells (middle panel) and successful restoration of the canonical sequence following CRISPR correction (lower panel). The designed silent PAM mutation successfully introduced by CRISPR is highlighted (see methods). **(C)** qRT-PCR demonstrating increased *LMNA* expression in cardiac organoids following correction of the *LMNA* mutation. (n = 4 mutant, n = 6 corrected, p = 0.01). **(D-E)** Western blot quantification (n = 3) and representative blots for *LMNA* in control (SCTi003-A), *LMNA*-mutant, and *LMNA-*corrected cardiac organoids. Lamin A and Lamin C were normalized to β-actin for quantification (Lamin A p = 0.0656, Lamin C *p = 0.0148) **(F)** Isoform-level quantification of *LMNA* transcripts in *LMNA-*mutant cardiac organoids. TPM = Transcripts per million. **(G)** Schematic illustrating aberrant exon 6 splicing caused by the c.937-1G>A mutation compared with canonical splicing. **(H)** Quantification of *LMNA* isoforms with normal, and with an aberrantly spliced short, and long Exon 6 (% of total *LMNA* expression), in control (SCTi003-A) and *LMNA*-mutant cardiac organoids.

Long-read RNA sequencing revealed expression of multiple *LMNA* isoforms in cardiac organoids, including the predominantly expressed canonical lamin A and lamin C transcripts, along with several low-abundance, alternatively spliced variants (Figure 1F). Lamin C is generated from the same primary transcript as lamin A through alternative 3’-end processing, resulting in a distinct C-terminal sequence and the exclusion of the final two coding exons of lamin A. Notably, the shorter lamin C-encoding transcripts also consistently lacked three annotated upstream untranslated exons present in lamin A transcripts. Although this observation is unexpected, the robust expression of both lamin A and lamin C, together with the absence of mixed splice isoforms, suggests coordinated isoform-specific processing of *LMNA* transcripts in cardiac organoids. Aberrant splicing of exon 6 was detected in *LMNA*-mutant organoids, involving the use of two cryptic splice acceptor sites located 8 bp downstream and 8 bp upstream of the canonical site (Figure 1G), both resulting in premature stop codons at amino acid positions 476 (elongated) and 364 (truncated). Transcripts containing a truncated exon 6 accounted for approximately 1.7% of the total *LMNA* transcripts (transcripts per million, TPM), whereas those with an elongated exon 6 represented 2.1% (Figure 1H). Both aberrant isoforms were exclusively detected in *LMNA-* mutant organoids and were absent in the wildtype (Supplemental Figure 2A), as well as in human hearts (Supplemental Figure 2B). Together, these findings identify *LMNA* c.937-1G>A as a novel pathogenic splice-site variant that disrupts pre-mRNA processing, generating premature termination codon-containing transcripts that are consistent with degradation likely through nonsense-mediated mRNA decay (NMD). The low abundance of these aberrant transcripts, together with reduced *LMNA* mRNA and protein levels and their restoration upon genetic correction, highlights c.937-1G>A as a mutation causing *LMNA* haploinsufficiency.

### *LMNA* Haploinsufficiency Drives Dysfunctional Calcium Dynamics and Arrhythmogenic Susceptibility

To assess calcium dynamics, we performed calcium imaging in whole cardiac organoids, as impaired calcium handling is a hallmark of DCM [10–12]. *LMNA*-mutant organoids exhibited a pronounced depression of systolic calcium levels compared with control organoids (Figure 2D). Interestingly, diastolic calcium levels were largely preserved in *LMNA*-mutant organoids (Figure 2E), suggesting a preferential impairment of systolic calcium release and calcium cycling kinetics rather than resting calcium homeostasis. Consistent with early calcium-handling abnormalities described in dilated cardiomyopathy, *LMNA*-mutant organoids demonstrated significantly lower amplitudes (Δ[Ca^2+^]_i_) (Figure 2F) and reduced upstroke and decay velocities (Figure 2 G,H), highlighting impaired calcium handling kinetics in *LMNA*-mutant organoids. In contrast, *LMNA*-corrected organoids showed substantial attenuation of these defects. Systolic [Ca^2+^]_i_ was significantly improved (Figure 2D*, p* < 0.05), as were calcium transient amplitudes (Δ[Ca^2+^]_i_) (Figure 2F*, p* = 0.004, 0.011, and 0.013 at increasing pacing rates). Upstroke velocity was also significantly improved to toward control levels (Figure 2G*, p* = 0.005, 0.011, and 0.016 at increasing pacing rates) while decay velocities trended toward control values (Figure 2H).

**Figure 2.**
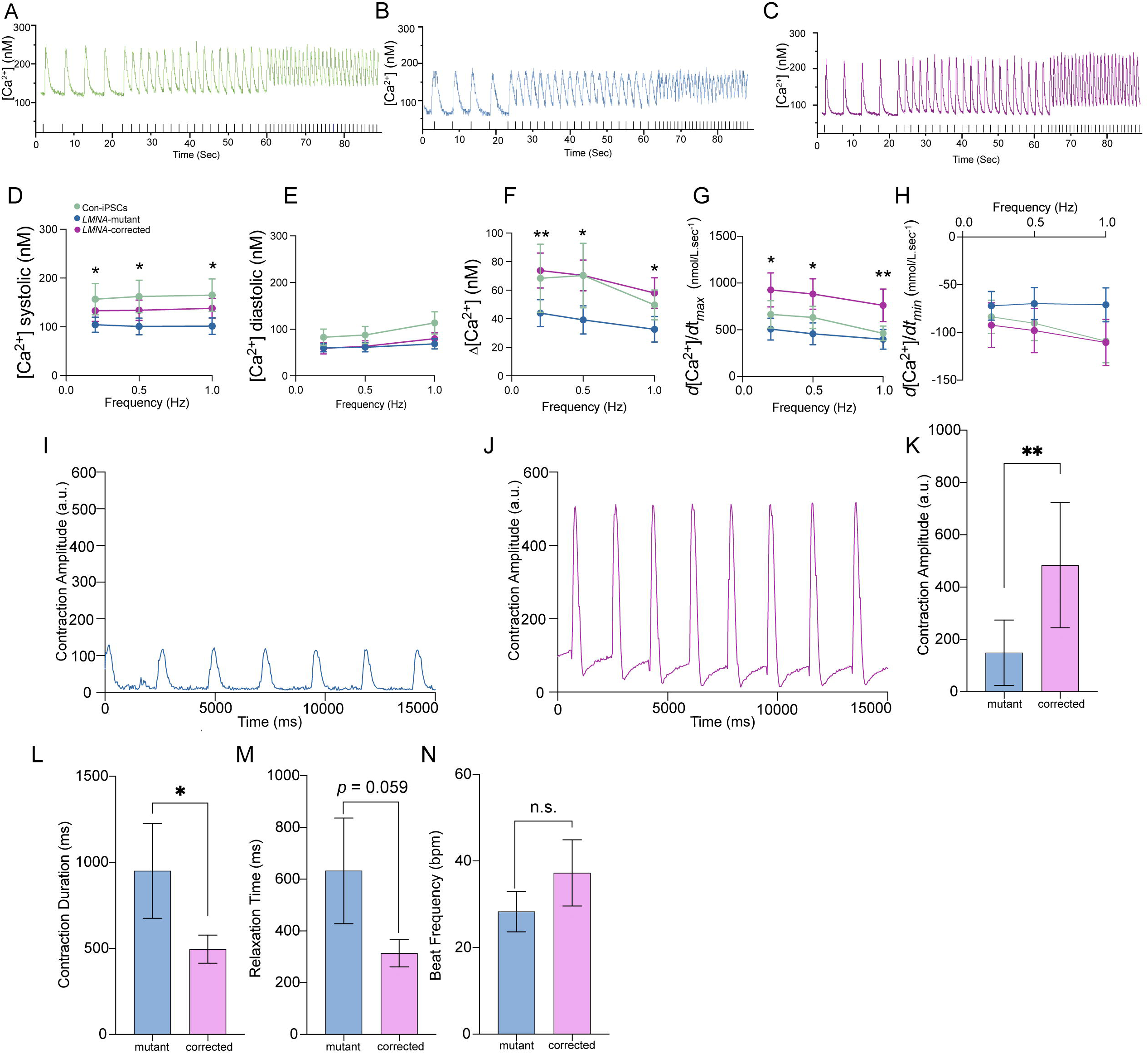
*LMNA*-mutant Organoids Demonstrate Calcium handling alterations. **(A-C)** Representative recorded traces of intracellular Ca^2+^ transients evoked by electric field stimulation at 0.2, 0.5 and 1 Hz in cardiac organoids from **(A)** Control (green), **(B)** *LMNA-*mutant (blue) and **(C)** *LMNA*-corrected (pink), respectively. (*LMNA*-corrected n = 15, *LMNA*-mutant n = 16) **(D)** Systolic Ca^2+^ levels in response to increasing pacing rate (*p* = 0.045, 0.041, and 0.035 at increasing pacing rates). **(E)** Diastolic Ca^2+^ levels in response to pacing **(F)** Ca^2+^ transient amplitude in response to pacing (p = 0.004, 0.011, and 0.013 at increasing pacing rates). **(G)** Maximal upstroke velocity *(d[Ca^2+^]/dt_max_)* at the different pacing rates (p = 0.005, 0.011, and 0.016 at increasing pacing rates). **(H)** Maximal decay velocity *(d[Ca^2+^]/dt_min_)* at the different pacing rates. **(I, J)** Representative contraction traces from *LMNA-mutant* and *LMNA*-corrected organoids (*LMNA*-corrected n = 6, *LMNA*-mutants n = 4). **(K)** Quantification of contraction amplitude (*p = 0.0379). **(L)** Contraction Duration (*p=0.0453). **(M)** Relaxation Time (ms) **(N)** Beat Frequency quantification (bpm). Data for control organoids (SCTi003-A) are shown in green, *LMNA*-mutant organoids in blue, and CRISPR-corrected organoids in pink. Data are represented as mean ± SEM.

*LMNA*-mutant organoids also exhibited a higher incidence of autonomous firing and arrhythmic events (Supplementary Figure 3 C,D), whereas *LMNA*-corrected organoids showed fewer arrhythmias and more closely resembled the independent control line, SCTi003-A). Although *LMNA*-mutant organoids displayed a higher proportion of spontaneous Ca^2+^ transients than controls, this trend did not reach statistical significance (Supplementary Figure 3A). Together, these findings complement the pacing-induced calcium defects observed in *LMNA*-mutant organoids and further establish impaired calcium handling as a direct consequence of reduced *LMNA* dosage.

### *LMNA-mutant* Cardiac Organoids Display Impaired Contractile Dynamics

To assess mechanical function, we quantified spontaneous contractile activity using quantitative optical motion tracking (Figure 2I-N). *LMNA-*mutant organoids exhibited significantly reduced contraction amplitudes compared to CRISPR-corrected organoids, which displayed uniform, high-amplitude rhythmic contractions (Figure 2J). Consistent with these observations, *LMNA*-mutant organoids showed a significant reduction in overall contraction amplitude relative to the isogenic CRISPR-corrected controls (Figure 2K, *p* < 0.01), shortened contraction duration (Figure 2L, *p* < 0.05), and a tendency toward increased relaxation times (Figure 2M, *p* = 0.05). In contrast, the frequency of spontaneous beating was not significantly altered by *LMNA* mutation (Figure 2N). Together, these results indicate that *LMNA* haploinsufficiency compromises cardiomyocyte contractile mechanics by reducing contractile force and altering contraction-relaxation kinetics.

### Cell-Type Specific Transcriptional Remodeling in *LMNA-mutant* Cardiac Organoids

To define early cell-type specific effects, we performed single-nucleus RNA-seq (snRNA-seq). UMAP projection revealed distinct clustering of various cardiac cell types, including atrial and ventricular cardiomyocytes, SAN pacemaker cells, fibroblasts, epicardial cells, vascular smooth muscle cells, AVN pacemaker cells, and endothelial cells (Figure 3A-C). Differential expression analysis of *LMNA*-mutant and CRISPR-corrected organoids revealed transcriptional changes in all cell types, however, epicardial and fibroblast cell types exhibited the greatest change in the number of significantly differentially expressed genes (DEGs) (FDR < 0.05), with 2,196 and 1,846 DEGs, respectively. Other cell types, including AVN pacemaker cells and atrial cardiomyocytes, demonstrated comparatively fewer transcriptional changes, with 88 and 134 DEGs, respectively (Figure 3B).

**Figure 3:**
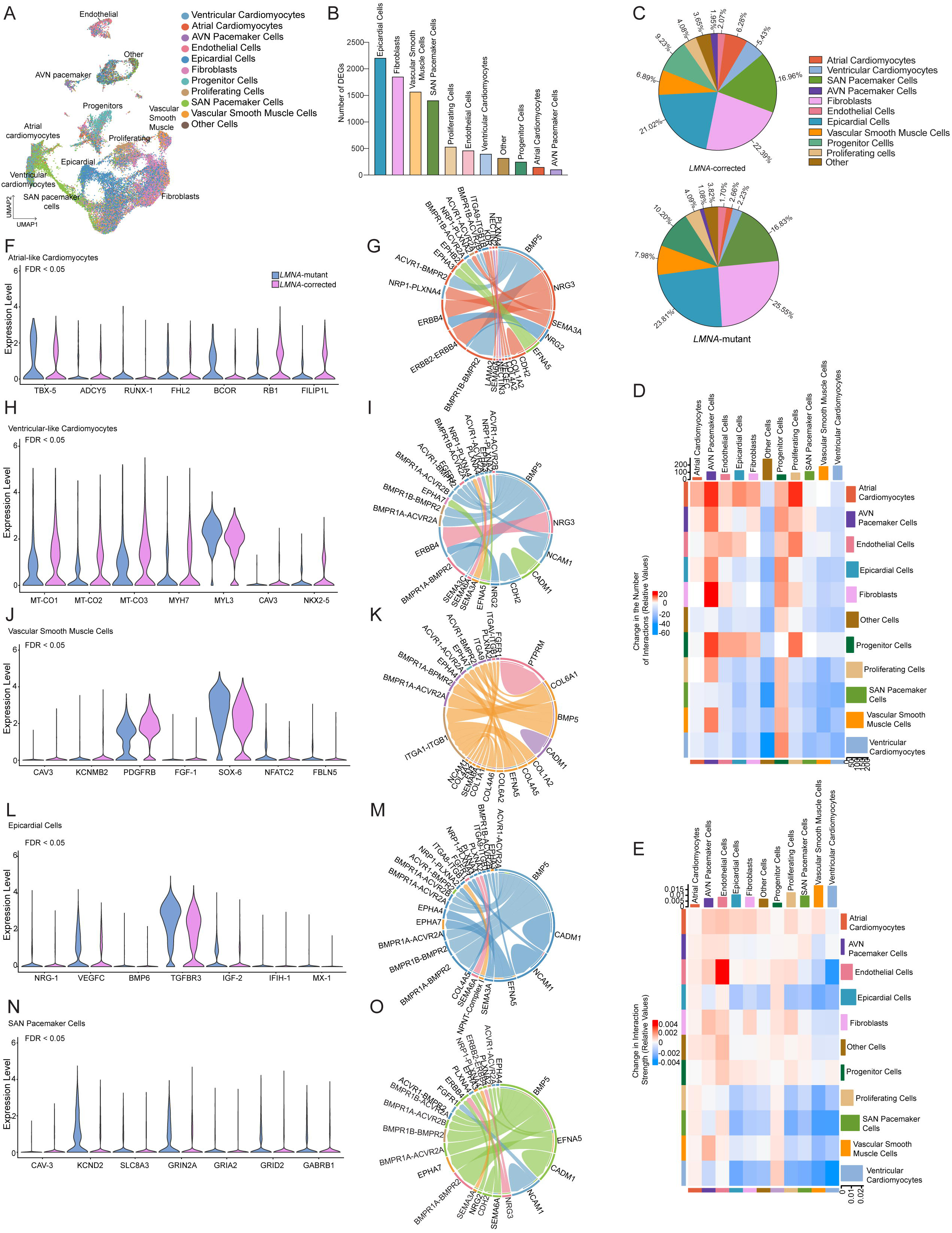
Single-nucleus Transcriptomics Reveals Lineage-Specific Remodeling Across *LMNA-mutant* Cardiac Organoids. **(A)** UMAP projection of Single-nucleus RNA sequencing data from *LMNA*-mutant and *LMNA*-corrected cardiac organoids, colored by annotated cell type. **(B)** Bar graph summarizing the number of significantly differentially expressed genes (DEGs; FDR < 0.05) identified in each major cardiac lineage comparing *LMNA*-mutant to corrected organoids. **(C)** Pie charts showing the relative cellular composition of *LMNA*-mutant and *LMNA*-corrected organoids. **(D)** Differential interaction number (count) heatmap derived from CellChat analysis, organized by signaling source (rows) and target (columns). Red boxes indicate increased number of interactions in *LMNA*-mutant organoids, whereas blue indicates decreased interactions relative to corrected organoids. **(E)** Differential interaction weight heatmap representing aggregate communication probability (signaling strength) between cell types. Red boxes indicate increased signaling strength in *LMNA*-mutant organoids, whereas blue indicates decreased signaling strength relative to corrected organoids. **(F-O)** Violin plots depicting differentially expressed genes (FDR < 0.01) for the indicated cell types, as well as chord diagrams depicting top unique 20 potential differential ligand-receptor interactions inferred by CellChat analysis. Chord width is proportional to the inferred interaction strength (communication probability) of ligand-receptor interactions. Expression values are shown on a log2-normalized scale.

Analysis of inferred intercellular communication networks revealed a redistribution of signaling activity across cardiac lineages in *LMNA*-mutant organoids. Atrial cardiomyocytes showed the largest increase in outgoing interactions, with an increase in signaling originating from endothelial and progenitor populations (Figure 3D). In contrast, SAN pacemaker cells and ventricular cardiomyocytes exhibited a broad reduction in outgoing interactions. Quantification of aggregate signaling strength further identified endothelial cells as the dominant source of amplified signaling, with additional increases observed in atrial cardiomyocytes and fibroblasts (Figure 3E). For endothelial cells, inferred increase in homophilic PTPRM-PTPRM interactions were the main contributor to this increased signaling change (Supplemental Figure 4A). PTPRM is a receptor tyrosine phosphatase that mediates homophilic cell-cell adhesion and regulates cadherin-dependent junctional signaling, suggesting altered endothelial cell-cell adhesion and junction associated signaling in *LMNA*-mutant organoids [13].

Despite substantial heterogeneity in transcriptional remodeling across cell types, three intercellular signaling axes were consistently enriched across cell types, indicating a shared component in the response to *LMNA* haploinsufficiency (Figure 3G-O). First, extracellular matrix (ECM)-integrin signaling was the most pervasive differential pathway and was enriched across all six analyzed cell types, including atrial and ventricular cardiomyocytes, vascular smooth muscle cells, epicardial cells, sinoatrial node (SAN) pacemaker cells, and fibroblasts. Collagen-integrin interactions involving *COL1A1, COL1A2, COL3A1,* and *COL4A1* engaging ITGA1/ITGA2-ITGB1 receptors were repeatedly identified as the strongest differential ligand-receptor pairs, indicating a broadly increased ECM-integrin signaling in *LMNA*-mutant organoids. Second, BMP family morphogen signaling, predominantly mediated by BMP5, was enriched across five of the six cardiac lineages, including atrial and ventricular cardiomyocytes, vascular smooth muscle cells, epicardial cells, and SAN pacemaker cells, but was not prominently represented among the top differential interactions in fibroblasts. *BMP5*-associated interactions involving multiple activin/BMP receptor combinations, including BMPR1A/B-BMPR2 and ACVR1-BMPR2, indicate a broad activation of morphogen signaling classically associated with cardiac remodeling and stress adaptation. Finally, adhesion and guidance-associated signaling pathways, including ephrin (EPHA-EFNA), cadherin-mediated adhesion, and semaphorin signaling, were enriched across all six cell types, although with lineage-specific ligand-receptor patterns. These three pathways are well established in cardiac development and injury responses, and their coordinated enrichment indicates that *LMNA* haploinsufficiency induces intercellular communication programs associated with cardiac remodeling and injury at the earliest stages of disease, as represented by the cardiac organoid model, even in the absence of overt damage or hemodynamic stress. Cell-type-specific deviations from this shared framework are described in the following sections. All reported differentially expressed genes met the significance threshold of FDR < 0.05. Although some transcriptional changes were modest in magnitude, they occurred in distinct subsets of cells, consistent with the heterogeneous expression patterns shown in Supplementary Figure 4.

#### Atrial-like Cardiomyocytes

Within the atrial-like cardiomyocyte population, *LMNA* mutations most notably affect the transcriptional programs governing atrial identity, calcium handling, and stress signaling (Figure 3F, G, Supplemental Figure 4F). Upregulated genes in mutant atrial-like cardiomyocytes included *TBX5*, a central regulator of atrial electrophysiology and conduction system maturation, and increased *ADCY5,* a key regulator of cAMP-dependent calcium signaling and β-adrenergic stress responses (Figure 3F) [14, 15]. Concurrently, mutant organoids also exhibited increased expression of injury-associated transcription factors *RUNX1* and *FHL2* [16, 17], together with increased expression of the chromatin regulator *BCOR*, indicating a broader remodeling of transcriptional regulatory programs. The reduced expression of genes associated with structural organization, cell cycle restraint, and cellular homeostasis, including *FILIP1L, RB1*, and APOA1, is consistent with altered cellular homeostasis and a stress-responsive phenotype. We further found enrichment of ERBB receptor-mediated signaling, with the strongest differential interactions including NRG3-ERBB2/ERBB4 and NRG3-ERBB4. In addition, lineage-specific guidance and adhesion interactions, including SEMA3A-NRP1-PLXNA4 and SEMA3A-NRP1-PLXNA2, homophilic CDH2-CDH2 adhesion, EFNA5-EPHB2 ephrin signaling, and COL1A2-ITGA9-ITGB1 ECM-integrin signaling were increased. Collectively, *LMNA*-mutant atrial-like cardiomyocytes exhibit injury-associated transcriptional activation and altered ERBB, guidance, adhesion, and ECM-associated signaling, consistent with a stress-responsive remodeling state.

#### Ventricular-like Cardiomyocytes

*LMNA-*mutant ventricular-like cardiomyocytes revealed transcriptional alterations affecting genes involved in mitochondrial energy metabolism, contractile architecture, and cardiomyocyte maturation. Specifically, the suppression of mitochondrial respiratory chain components *MT-CO1, MT-CO2, and MT-CO3* (Figure 3H,I, Supplemental Figure 4G), *MT-CYB, MT-ND1, MT-ND3,* and *NDUFB9* (Supplemental Figure 4A) is consistent with impaired expression of oxidative phosphorylation machinery. This was accompanied by downregulation of the contractile and membrane-structural genes *MYH7, MYL7,* and *CAV3* and reduced expression of *NKX2-5,* a transcriptional regulator essential for ventricular lineage specification and maturation, together supporting impaired maturation [18, 19]. Additionally, we found upregulated genes involved in adhesion and remodeling, including *EPHA3* and *EPHA7* (Supplemental Figure 4A), which have defined roles in cardiac patterning and septal and valvular morphogenesis [20]. Further, increased? ERBB receptor signaling, with the strongest differential interaction being NRG3-ERBB4, potentially arising from endothelial cells, indicates increased influence of endothelial-derived paracrine signaling. *LMNA*-mutant ventricular cardiomyocytes also demonstrated enhanced cell-cell adhesion and connectivity, including homophilic CDH2-CDH2 interactions, which have been implicated as contributors to cardiac remodeling and regenerative responses following injury [21]. In addition, we found NCAM1-NCAM1 signaling, consistent with the upregulation of NCAM1 in failing cardiomyocytes [22], as well as EFNA5-EPHA7 ephrin interactions potentially originating from ventricular and SAN pacemaker populations. Taken together, these transcriptional and intercellular signaling alterations define a ventricular-specific remodeling program marked by impaired metabolic and contractile maturation, altered ERBB-mediated communication, reinforced cell-cell adhesion, and an activation of stress-associated signaling in *LMNA*-mutant organoids.

#### Vascular Smooth Muscle Cells

*LMNA-*mutant vascular smooth muscle cells (VSMCs) displayed coordinated transcriptional changes indicative of impaired contractile homeostasis and an increased stress-responsive remodeling phenotype. Notably, we found marked reductions in expression of *CAV3* and *KCNMB2* (Figure 3J, Supplemental Figure 4G), genes that contribute to maintaining contractile and ion-handling programs in VSMCs [23, 24]. Additionally, mutant VSMCs showed reduced expression of key ciliary and trafficking regulators, including *CC2D2A, CC2D2B,* and *MAK* (Supplemental Figure 4B), supporting alterations in primary cilium-associated and mechanosensory signaling programs [25]. The expression of key growth factor signaling mediators *PDGFRB, FGF1,* and VEGFA was also decreased, suggesting an attenuation of pathways implicated in vascular homeostasis [26, 27]. In parallel, upregulated stress- and injury-response genes, including *LPAR1, OLR1, IL6R,* and *AGTR1* (Supplemental Figure 4B) [28–30], along with remodeling-associated transcriptional regulators, such as SOX6 and NFATC2. This expression pattern is consistent with the early activation of stress-responsive VSMC programs [31]. Further, ECM deposition and remodeling genes, including *COL5A1, COL11A1, FBLN5*, and *NPNT* (Supplemental Figure 4B), were also increased, supporting a shift toward a fibroblast-like transcriptional state, a phenotypic switch reported during vascular injury and remodeling [32]. Increased expression of non-canonical transcription factors, such as *SOX5* and *MITF*, further indicates the remodeling of VSMC transcriptional identity in the context of *LMNA* haploinsufficiency. Consistent with these signaling changes, *LMNA*-mutant VSMCs also demonstrated differential interactions, reflecting altered structural and guidance signaling, including homophilic CDH2-CDH2 adhesion, and enhanced EPHA-EFNA ephrin signaling (Figure 3K), collectively defining a stress-responsive, matrix remodeling VSMC communication state.

#### Epicardial Cells

*LMNA-*mutant epicardial cells showed upregulation of genes associated with epicardial injury response, including *NRG1, VEGFC, BMP6, TGFBR3*, and *IGF2* (Figure 3L, Supplemental Figure 4H) [33–35]. In addition, canonical interferon-stimulated genes (ISGs), including *IFIH1, MX1*, and IFI44L, along with additional ISGs such as *SP100* (Supplemental Figure 4C), were upregulated, consistent with the activation of interferon-associated signaling in cardiac-associated injury responses [36]. Given the established role of IFIH1/MDA5 in protecting against myocardial injury, ISG induction reflects the upregulation of an injury-responsive interferon program [37]. Epicardial cells also displayed enhanced laminin-integrin signaling and EPHA-EFNA guidance interactions (Figure 3M), consistent with altered adhesion. Together, these findings indicate that *LMNA*-mutant epicardial cells adopt an injury-responsive, remodeling-associated state characterized by interferon activation and altered adhesion- and guidance-mediated paracrine communication.

#### SAN Pacemaker Cells

*LMNA*-mutant SAN pacemaker cells demonstrated altered expression of pacemaker-associated genes and increased expression of neuronal signaling-associated genes, suggesting a partial shift toward a more neuronal-like transcriptional program. *CAV3* expression was markedly reduced in *LMNA-*mutants (Figure 3N, Supplemental Figure 4I), which encodes a key structural regulator that promotes the clustering of pacemaker ion-channel complexes, including HCN4, Cav1.3, and Cav3.1 [38]. In parallel, *LMNA*-mutants displayed increased expression of SAN-associated calcium and potassium channel genes *KCND2* and *SLC8A3/NCX3* (Figure 3N), the latter encoding a Na^+^/Ca^2+^ exchanger involved in Ca^2+^ homeostasis. Interestingly, we found increased expression of glutamatergic and GABAergic receptor genes, such as *GRIN2A, GRIA2*, *GRID2*, and *GABRB1*. Because SAN pacemaker cells share transcriptional features with neurons [39], these changes suggest a shift toward increased neuronal-like signaling features. SAN pacemaker cells exhibited a distinct repertoire of non-shared interactions, including CADM-mediated homophilic adhesion, NCAM1-NCAM1 signaling, SEMA6-plexin guidance interactions, FN1-integrin signaling, and EPHA-EFNA ephrin interactions (Figure 3O). Collectively, these data suggest that *LMNA* haploinsufficiency drives SAN pacemaker cells toward a neuronal-like transcriptional state, coupled with impaired ion channel organization.

#### Fibroblasts

*LMNA*-mutant fibroblasts exhibited transcriptional changes in extracellular matrix organization, inflammatory signaling, and regulatory pathways associated with fibroblast activation. Genes involved in extracellular matrix remodeling and cytoskeletal organization, including *ARHGAP44, ARHGAP24, DOCK5, FRZB, FBLN5, PDPN,* and *COL24A1* (Figure 4A, B), were upregulated, along with increased expression of inflammatory and stress-associated genes such as *CD44, IL6R, TNFAIP3, EPSTI1,* and *AGTR1* (Supplemental Figure 4D). Notably, *CD44*, a key mediator of fibroblast migration, matrix remodeling, and inflammatory signaling in the heart, was robustly induced, consistent with an activated fibroblast state [40]. In parallel, *LMNA*-mutant fibroblasts upregulate genes implicated in stress-adaptive and modulatory signaling, including *BMP6* and *NPNT* [41]. *BMP6*, which has been reported to exert protective and antifibrotic effects during cardiac remodeling, was expressed at higher levels [42]. *PDZK1*, which encodes a scaffolding protein implicated in restraining cardiac fibroblast activation, was downregulated [43]. In addition, *MMP24* and *ASPN* (Supplemental Figure 4D), two regulators of extracellular matrix turnover and TGF-β-associated matrix control that have been implicated in limiting pathological fibrosis, were reduced [44]. *LMNA*-mutant fibroblasts further demonstrated enhanced laminin-integrin and FN1-associated signaling, along with EPHA-EFNA ephrin interactions and CADM- and NCAM-mediated adhesion, reflecting reinforced matrix engagement and altered intercellular connectivity (Figure 4C). Together, these findings indicate that *LMNA*-mutant fibroblasts adopt an activated, profibrotic transcriptional state characterized by coordinated extracellular matrix remodeling, inflammatory signaling, and altered intercellular connectivity.

**Figure 4.**
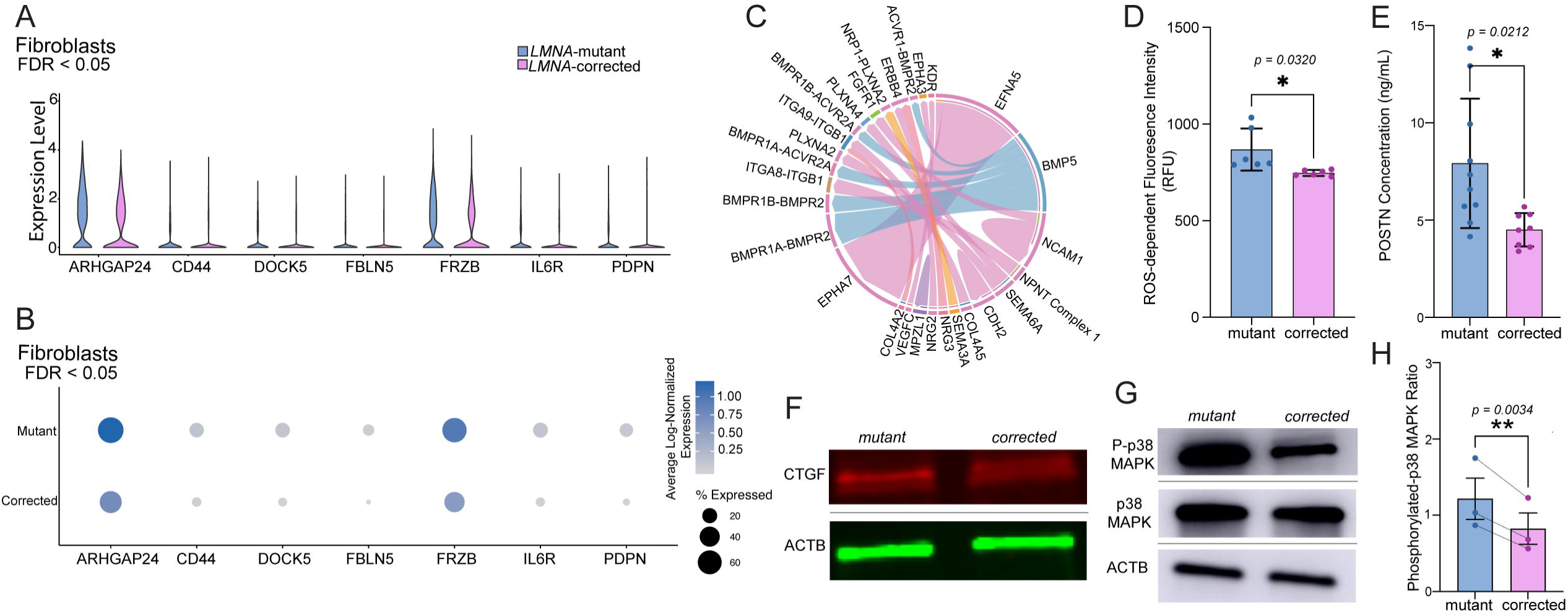
*LMNA*-mutant cardiac organoids exhibit fibroblast activation, oxidative stress. **(A)** Violin plots showing increased expression of fibroblast activation and extracellular matrix-associated genes in fibroblasts from *LMNA-mutant* compared with corrected organoids. **(B)** Dot plot displaying average expression and % of fibroblasts expressing the respective genes. **(C)** Chord diagram depicting differential ligand-receptor signaling involving fibroblasts identified by CellChat analysis. **(D)** Quantification of reactive oxygen species (ROS) dependent fluorescence intensity levels showing elevated oxidative stress in *LMNA*-mutant organoids relative to isogenic corrected controls (*\*p* = 0.0320). **(E)** Quantification of periostin (*POSTN*) secretion in conditioned media from *LMNA*-mutant and corrected organoids, demonstrating increased fibrotic output in Mutants compared with corrected organoids (*\*p* < 0.05). **(F)** Western blot analysis of *CTGF* protein expression in *LMNA*-mutant and corrected organoids, with *ACTB* as a loading control. **(G)** Western blot analysis of *p38 MAPK* and *P-p38 MAPK* protein expression in *LMNA*-mutant and corrected organoids, with *ACTB* as a loading control. **(H)** Quantification of the phosphorylated *p38 MAPK/total p38 MAPK* ratio in *LMNA*-mutant and *LMNA*-corrected cardiac organoids. Data are presented as mean ± SEM with individual biological replicates shown and paired by lines (*n* = 3). Statistical significance was determined using a ratio paired *t*-test (\*\**p* = 0.0034).

### *LMNA-mutant* Cardiac Organoids Exhibit Fibroblast Activation and Oxidative stress

To assess a potential early fibrotic phenotype, we assessed reactive oxygen species (ROS) and fibroblast activation by quantifying periostin (*POSTN*) secretion, a marker of activated fibroblasts, and matricellular protein induced during cardiac injury and fibrotic remodeling [45]. *LMNA*-mutant organoids exhibited higher levels of ROS (*p* < 0.05, Figure 4D) and secreted higher levels of *POSTN* than *LMNA*-corrected organoids (*p* < 0.05, Figure 4E). We further evaluated connective tissue growth factor (*CTGF*) protein levels, a key downstream mediator of fibrosis, by western blotting [46]. *LMNA*-mutant organoids exhibited increased *CTGF* protein levels compared to isogenic corrected controls (*n = 3*, Figure 4F), further supporting activation of profibrotic signaling in the mutant condition. To further assess the activation of fibrosis-associated signaling, we evaluated *p38-MAPK* phosphorylation by western blotting (Figure 4 G,H). While total p38 levels were comparable between groups, phosphorylated *p38-MAPK* (p-p38)/total *p-38-MAPK* ratio was increased in *LMNA*-mutant organoids (*p* = 0.0034, Figure 4H), consistent with enhanced activation of MAPK signaling in the mutant condition. Together, these findings demonstrate that *LMNA*-mutant cardiac organoids exhibit an early profibrotic phenotype characterized by increased oxidative stress, fibroblast activation, and activation of fibrosis-associated signaling pathways.

### Antifibrotic treatment attenuates profibrotic signaling in *LMNA*-mutant cardiac organoids

To determine whether the early fibrotic signaling observed in *LMNA*-mutant organoids is therapeutically targetable, we treated *LMNA*-mutant organoids with the antifibrotic agents nintedanib and pirfenidone, both of which have demonstrated antifibrotic efficacy and have been explored for cardioprotective effects [47, 48]. Nintedanib treatment significantly reduced the secretion of periostin (POSTN) compared with vehicle-treated controls (Figure 5A, *p* < 0.05), whereas pirfenidone-treated organoids did not exhibit a significant reduction in POSTN secretion. In contrast, both nintedanib and pirfenidone reduced *CTGF* protein levels (Figure 5 B, C), indicating a partial attenuation of profibrotic signaling.

**Figure 5.**
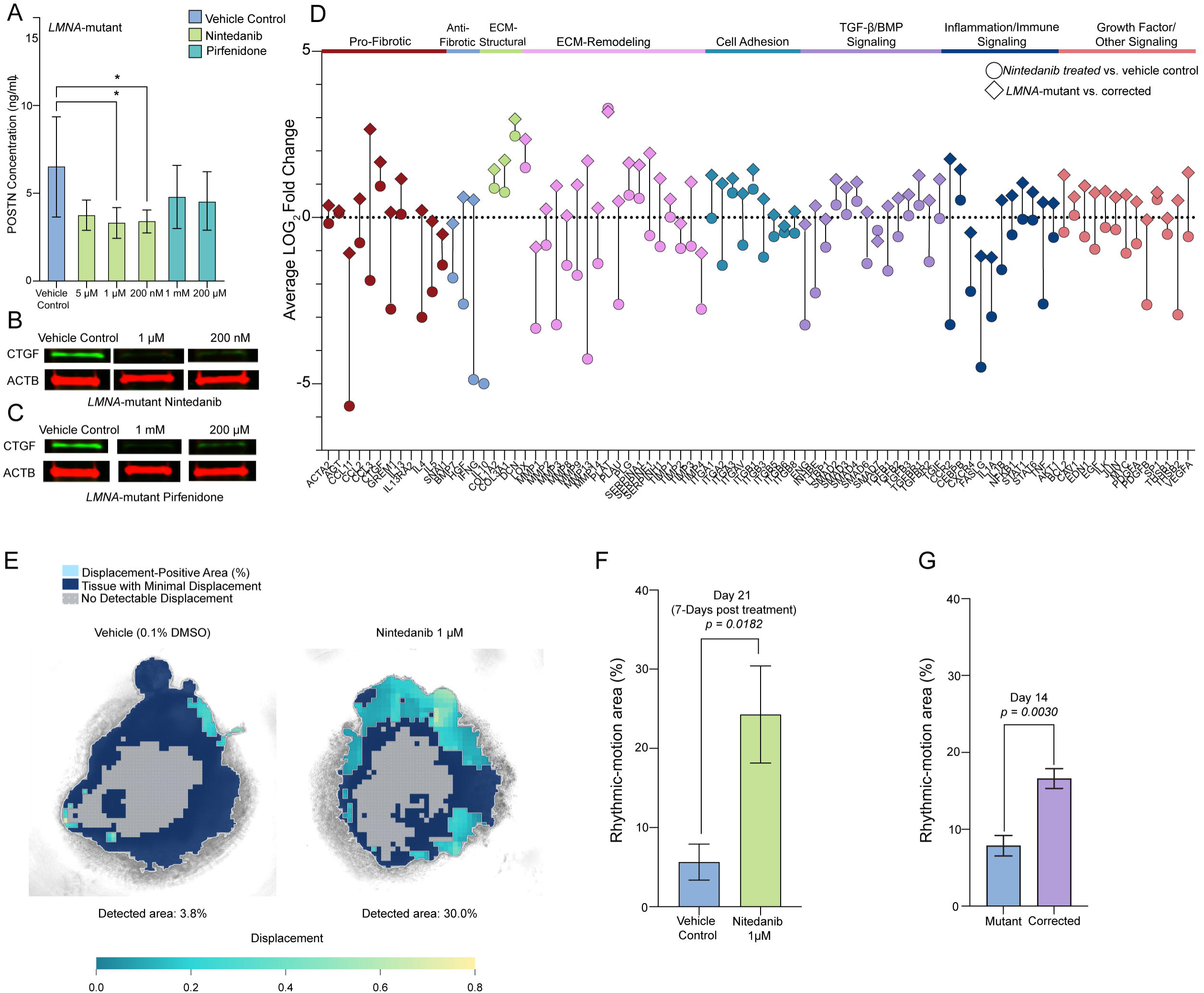
Antifibrotic treatment attenuates profibrotic signaling in *LMNA*-mutant cardiac organoids. **(A)** Quantification of periostin *(POSTN)* secretion in conditioned media from L*MNA*-mutant cardiac organoids treated with nintedanib (5 µM, 1 µM, 200 nM) or pirfenidone (1 mM, 200 µM), showing reduced *POSTN* secretion following nintedanib treatment compared with vehicle control (*p* < 0.05). **(B)** Western blot analysis of CTGF protein levels in *LMNA*-mutant organoids treated with nintedanib (1 µM and 5 µM). **(C)** Western blot analysis of *CTGF* protein levels in *LMNA*-mutant organoids treated with pirfenidone (1 mM and 200 µM). (**D**) RT^2^ Profiler PCR array analysis of fibrosis-associated genes. Log_2_ fold changes are shown for nintedanib-treated versus vehicle-treated *LMNA*-mutant organoids (circles) and untreated *LMNA*-mutant versus *LMNA*-corrected organoids (diamonds). Genes are grouped according to functional categories, including pro-fibrotic, anti-fibrotic, extracellular matrix (ECM) structural and remodeling genes, cell adhesion, TGF-β/BMP signaling, inflammation/immune signaling, and growth factor/other signaling. The dotted line indicates no change (log_2_ fold change = 0). **(E)** Representative spatial displacement maps of vehicle-treated and nintedanib-treated (1 µM) following 7-day treatment. Rhythmic motion was assessed using particle image velocimetry (PIV)-based analysis, with displacement-positive regions shown in light blue, regions with minimal displacement in dark blue, and regions without detectable displacement in gray. The percentage of organoid area exhibiting rhythmic motion is indicated below each image. **(F)** Quantification of rhythmic-motion area in *LMNA*-mutant cardiac organoids following seven days of nintedanib treatment (1 µM), showing an increased proportion of tissue exhibiting rhythmic motion compared with vehicle control (*p* = 0.0182). **(G)** Quantification of rhythmic-motion area in untreated *LMNA*-mutant and isogenic-corrected cardiac organoids at day 14, showing increased rhythmic-motion area in corrected organoids compared with mutant organoids (*p* = 0.0030).

To further characterize the effects of nintedanib, we quantified the expression of 84 genes involved in fibrosis and tissue remodeling using a quantitative PCR array (Figure 5D). Several fibrosis-associated genes elevated in *LMNA*-mutant relative to isogenic-corrected organoids were reduced following nintedanib treatment, including *CCL3* (mutant/corrected log2FC = 2.65; nintedanib/vehicle log2FC = −1.89), *CCR2* (1.76; −3.22), *MMP13* (1.70; −4.25), *MMP3* (0.95; −3.22), *THBS2* (0.51; −2.92), and *TGFB1* (0.34; −1.60). These changes encompassed chemokine-associated inflammatory signaling (*CCL3* and *CCR2*), extracellular matrix remodeling and organization (*MMP3*, *MMP13*, and *THBS2), and the* expression of the profibrotic ligand *TGFB1*. The expression of structural collagen genes was not uniformly reduced. Together with reduced POSTN secretion and CTGF protein expression, these findings support an early pharmacologically responsive disease phenotype involving inflammatory and matrix-remodeling processes.

Although seven days of treatment with either pirfenidone or nintedanib did not significantly affect contraction amplitude, time to peak, peak-to-peak interval, or relaxation time (Supplemental Figure 4K-L), spatial motion analysis revealed a significant improvement in the extent of rhythmic contraction following treatment with 1 µM nintedanib. Specifically, the proportion of organoid area exhibiting rhythmic motion increased from 5.6% in vehicle-treated controls to 24.3% in nintedanib-treated organoids (*p* = 0.01; Figure 5). These findings suggest that nintedanib may improve the distribution of contractile activity without altering individual contraction kinetics, highlighting the potential for functional improvement during early *LMNA*-associated cardiac remodeling.

Together, these findings demonstrate that nintedanib attenuates profibrotic and inflammatory signaling while increasing the proportion of tissue exhibiting rhythmic contractile activity. These results suggest that early *LMNA*-associated cardiac remodeling is pharmacologically responsive at both the molecular and functional levels.

## DISCUSSION

*LMNA*-associated dilated cardiomyopathy (*LMNA*-DCM) is one of the most penetrant inherited cardiomyopathies, yet the early cellular events that initiate the disease remain poorly understood. Using patient-derived cardiac organoids, we identified early activation of multicellular remodeling and profibrotic signaling programs as key consequences of *LMNA* haploinsufficiency in self-patterning human cardiac organoids. Most mechanistic insights have come from murine models and two-dimensional iPSC-derived cardiomyocytes, which have been instrumental in defining downstream phenotypes but are limited in their ability to resolve early human-specific mechanisms and multicellular processes that drive disease onset.

In this study, we used an iPSC-derived human cardiac organoid model to characterize a novel *LMNA* mutation (c.937-1G>A), which we found leads to *LMNA* haploinsufficiency. We show that *LMNA* haploinsufficiency disrupts transcriptional programs across all major cardiac lineages and produces convergent remodeling of intercellular communication. This remodeling is characterized by prominent extracellular matrix-integrin interactions, recurrent engagement of BMP/TGF-β family signaling, and altered adhesion and guidance cues. Similar communication programs have been described in the context of cardiac injury and remodeling, including pressure overload, myocardial infarction, and fibrotic heart failure [49]. Notably, these pathways emerged in the absence of external insults typically associated with cardiac remodeling in human disease, such as ischemic injury, pressure overload, or myocardial infarction, suggesting that *LMNA* haploinsufficiency can initiate an early injury-associated signaling state. While these shared alterations were observed across cardiomyocytes, vascular smooth muscle cells, epicardial cells, SAN pacemaker cells, and fibroblasts, each lineage also exhibited signaling changes that were consistent with its functional role. These include ERBB signaling in cardiomyocytes, which regulates cardiomyocyte survival and stress adaptation, matrix-dominant programs in fibroblasts and VSMCs, reflecting their roles in extracellular matrix production and structural remodeling, and adhesion- and guidance-associated signaling in pacemaker and epicardial cells, consistent with their roles in tissue organization and intercellular communication. Together, these findings support a model in which *LMNA* haploinsufficiency promotes coordinated multicellular remodeling of cardiac intercellular communication and lineage-specific transcriptional adaptations.

*LMNA*-mutant organoids exhibited impaired intracellular calcium handling characterized by reduced systolic Ca^2+^ levels, diminished transient amplitude, and slowed calcium kinetics. Notably, diastolic Ca^2+^ levels were not significantly altered. These findings indicate impaired calcium cycling during excitation-contraction coupling rather than a generalized disruption of resting calcium homeostasis, and are consistent with early features of dilated cardiomyopathy, where impaired systolic Ca^2+^ transients and slowed Ca^2+^ kinetics arise from reduced sarcoplasmic reticulum Ca²^+^ release and reuptake before overt diastolic Ca²^+^ dysregulation develops [50]. *LMNA*-mutant organoids also showed a trend toward increased arrhythmic events, consistent with the electrophysiological instability frequently observed in *LMNA*-associated cardiomyopathy [51].

Fibroblasts in *LMNA*-mutant organoids underwent extensive transcriptional remodeling involving extracellular matrix organization and inflammatory signaling. Increased POSTN secretion, ROS, p38 MAPK phosphorylation, and CTGF protein levels, together with broader fibrosis-associated transcriptional changes, highlight the emergence of a fibrotic state. Antifibrotic treatment partially attenuated this phenotype. Nintedanib reduced POSTN secretion and CTGF levels, opposed selected fibrosis-associated transcriptional changes, and increased the proportion of organoid area exhibiting rhythmic motion. Neither nintedanib nor pirfenidone detectably improved contraction amplitude or timing over seven days. This response suggests that fibrotic/inflammatory signaling and matrix remodeling cause tissue-level impairments that can be targeted pharmacologically, beyond the direct effects of *LMNA* deficiency in cardiomyocytes.

In summary, this study demonstrates that *LMNA* haploinsufficiency initiates early and coordinated remodeling across multiple cardiac lineages, integrating electrophysiological instability with fibrotic and stress-responsive signaling pathways. By capturing these alterations in a human iPSC-derived cardiac organoid model, we identified multicellular disease mechanisms that precede overt cardiomyopathy and are difficult to resolve in conventional experimental systems. The ability of this platform to recapitulate key features of *LMNA*-associated cardiomyopathy, including calcium handling defects, arrhythmogenic susceptibility, and early fibroblast activation, within a scalable human tissue context, provides a framework for studying the early stages of disease pathogenesis. Collectively, these findings identify early multicellular remodeling and profibrotic signaling as a defining consequence of *LMNA* haploinsufficiency and support human cardiac organoids as a platform for investigating disease initiation and identifying therapeutic strategies capable of targeting early disease-associated signaling before overt structural pathology develops.

## METHODS

### Generation of Patient-Specific iPSCs

Somatic reprogramming was used to generate iPSC lines from peripheral blood mononuclear cells using the CytoTune-iPS 2.0 Sendai Reprogramming Kit (Invitrogen, A16517) following the manufacturer’s instructions.

### iPSC Culture and Maintenance

Control iPSCs (SCTi003-A, StemCell Technologies), *LMNA*-mutant, and corrected iPSCs were cultured feeder-free in supplemented mTESR plus media (#100-0276, StemCell Technologies) on a Matrigel Basement Membrane (#CB-40234A, Corning) diluted in DMEM/F-12 (#10-092-CV, Corning) at 37°C with 5% CO_2_. The media was replaced daily, and cells were passaged every 4-5 days using ReLeSR (#100-0483, StemCell Technologies) and resuspended in supplemented mTESR plus media. The cells were monitored for signs of spontaneous differentiation and routinely tested for confirmation of pluripotency markers SOX2, SSEA4, and OCT-4 (Supplementary Table 1) via immunocytochemistry before experimentation (Supplementary Figure 1). To confirm the pluripotency and differentiation capacity of the iPSCs, karyotyping and trilineage differentiation was performed by directing cells to differentiate into ectoderm, mesoderm, and endoderm lineages, with successful differentiation validated by expression of lineage-specific markers (Supplementary Table 2). Chromosome analysis by G-banding was completed using *LMNA*-mutant and corrected iPSCs at passage 35 by the WiCell Research Institute. At least 20 metaphase nuclei were counted, eight were analyzed, and four were karyogrammed at the 400-500 band level.

### Trilineage Differentiation and Immunostaining of Germ Layer Markers

Differentiation of iPSCs into ectoderm, mesoderm, and endoderm (Supplementary Figure) was performed using a STEMDiff Trilineage Differentiation Kit (#05230, StemCell TECHNOLOGIES) according to the manufacturer’s instructions. Immunostaining confirmed the expression of markers unique to each germ layer (Supplementary Figure). For immunocytochemistry, the cultures were fixed with 4% formaldehyde for 15 min. Fixed cells were washed three times with PBS for 5 min, and then incubated for 45 min in permeabilization/blocking (P/B) buffer comprised of PBS, 10% normal donkey serum (NDS), 0.3% Triton X-100, 0.05% Tween-20, and 0.3 M glycine. Primary antibodies were diluted in antibody dilution (ABD) buffer containing phosphate-buffered saline (PBS), 2% normal donkey serum (NDS), 0.1% Triton X-100, 0.05% Tween-20, and 0.02% sodium azide. After removing the P/B buffer, diluted primary antibodies were applied to the cells and incubated overnight at 4°C. The next day, the primary antibodies were removed, and the cells were washed three times with PBS for 5 min. Secondary antibodies were diluted 1:1000 in ABD buffer, added to the cells, and incubated for 1 h at room temperature in the dark. The cells were then washed three times with PBS for 5 min and incubated for 15 min with NucBlue (DAPI; Thermo Fisher Scientific, R37606). The NucBlue DAPI solution was replaced with PBS, and the cells were imaged using a Keyence BZ-X810 microscope.

#### CRISPR/Cas9 gene-editing

A chemically modified sgRNA (Synthego) and single-stranded oligo donor (ssODN) containing phosphorothioate bonds at the 5′ and 3′ ends (IDT Ultramer) were designed using CRISPOR (https://crispor.gi.ucsc.edu/) and the Benchling CRISPR design tools (https://www.benchling.com/crispr) (Supplemental Table 3). A silent PAM mutation was introduced to prevent Cas9 from re-cleaving. Human induced pluripotent stem cells (iPSCs) were maintained on vitronectin-coated plates in mTeSR Plus medium (#100-0276, STEMCELL Technologies) and passaged to reach 60-80% confluency on the day of editing.

Genome editing was performed by nucleofecting 200,000 iPSCs with Cas9 (1.6 µM)-sgRNA (5.6 µM) ribonucleoprotein (RNP) complexes and 3 µM ssODN using the Lonza P3 nucleofection solution (25 µL reaction volume) on a Lonza Amaxa 4D-Nucleofector (program CA-137). The nucleofected pool was seeded into a vitronectin-coated 12-well plate in mTeSR Plus containing 10 *μ*M Y-27632 and 2 *μ*M Thiazovivin (#S1459, Selleck Chemicals), with both ROCK inhibitors removed the following day. The cells were subsequently maintained with daily media changes until cryopreservation and genomic analysis. To estimate the knock-in efficiency, genomic DNA was extracted from the edited pool using QuickExtract (Lucigen), followed by Sanger sequencing and analysis using DECODR software. For clonal isolation, the edited iPSC pool was dissociated with Accutase (#07920, STEMCELL Technologies) for 10 min at 37°C, diluted to ∼1 cell/100 µL, and plated into rhLaminin-521-coated 96-well plates (Thermo Fisher) in mTeSR Plus supplemented with CloneR2 (#100-0691, STEMCELL Technologies).

After approximately two weeks, surviving colonies were dissociated with 50 µL Accutase for 5 min. Half of each well was transferred into tubes containing 200 *µ*L CryoStor CS10 freezing medium, sealed with push caps, and stored at -80°C. The remaining cells were used for genotyping. Genomic DNA was extracted using QuickExtract, and genomic regions flanking the *LMNA* correction site and the top five predicted off-target loci were PCR-amplified using Q5 Hot-Start High-Fidelity Polymerase (#M0493S, NEB) and primers containing M13 tails (Supplemental Table 2). Amplicons were submitted to GeneWiz for Sanger sequencing using M13-forward and M13-reverse primers (Supplemental Table 2).

### Cardiac Organoid Differentiation

*LMNA*-mutant and corrected iPSCs were differentiated into cardiac organoids as previously described [8], with a few minor changes: iPSCs were cultured in mTESR plus supplement media (#100-0276, StemCell Technologies) instead of Essential-8 media. Additionally, on day -2, 15,000 iPSCs were seeded per well on the Sarstedt Biofloat round base plates (#83.3925.400, Starstedt), rather than the Costar plates utilized in the protocol. For *LMNA-mutant* organoids CHIR concentration was also altered to enhance differentiation efficacy, ensuring comparability with wildtype organoids and minimizing cell line-specific differences; first dose of GSK3β inhibitor, CHIR-99021 (#SML1046, Sigma) was increased to a final concentration of 8 *μ*M and second dose of CHIR-99021 4 *μ*M. All differentiation conditions for Control-iPSCs were performed exactly as described in the published protocol, without modification.

### Long-Read RNA-sequencing

Full-length RNA isoform sequencing was performed by the Hussman Institute for Human Genomics (HIHG) Sequencing Core facility at the University of Miami using the PacBio (platform). High-quality total RNA was extracted from cardiac organoids, and cDNA synthesis and library preparation were carried out using the Iso-Seq express workflow (Pacific Biosciences) according to the manufacturer’s instructions. Reads were aligned to the human reference genome (GRCh38) using minimap2 (v2.28) with splice aware parameters. PacBio classification outputs were filtered to include *LMNA*-associated transcripts with TPM >0.5. Filtered GFF entries corresponding to expressed isoforms were visualized with ggtranscript using the functions *geom_range* and *geom_intron* to display exon-intron structures and UTRs. Transcripts were plotted using ggplot2 and final figures were exported as scalable vector graphics (SVG) for figure assembly.

### Western Blotting

Protein lysates were prepared from cardiac organoid cell pellets using RIPA lysis buffer supplemented with protease inhibitor tablets (#05892970001, Roche) and 15U of DNAse I (#E1010, ZymoResearch). Samples were incubated for 10 min at 37°C, sonicated (15 s x 3) and centrifuged at 15,000 x g for 15 min at 4°C for debris removal. Protein quantification was determined using BCA protein assay kit (#23209, Thermo Scientific). Equal amounts of protein were loaded for each sample (20 µg) mixed with 4x Laemmli sample buffer containing *β*-mercaptoethanol and heated to 95°C for 5 min before loading. Proteins were separated by SDS-Page on Mini-Protean TGX Stain-Free Precast Gels (#4561093, Bio-Rad) and transferred onto PVDF membranes (#1704156, BioRad). Membranes were blocked at 1 h at room temperature in 5% BSA in TBS-T buffer and then incubated overnight at 4°C with primary antibodies diluted in blocking buffer (Supplemental Table 1). After washing three times with TBS-T, membranes were incubated with LI-COR infrared dye-conjugated secondary antibodies. Bands were visualized using the LI-COR Odyssey imaging system and band intensities were quantified using the LI-COR Image Studio, normalized to β-actin (Supplemental Table 1) as a loading control.

### Quantitative Real-Time PCR (qPCR)

Total RNA was extracted from cardiac organoids using the Quick-RNA Microprep Kit (#R1051, Zymo Research) according to the manufacturer’s instructions. RNA concentrations were measured using the Denovix RNA quanitification Assay (#RNA-EVAL, DeNovix). cDNA synthesis was performed using the iScript cDNA synthesis Kit (Bio-Rad) in a 20 µl reaction volume. Quantitative PCR was carried out using the SYBR Green Master Mix (#1725271, Bio-Rad) on the Cielo qPCR system (Azure Biosystems). Each reaction contained 1 µL of cDNA template. Cycling conditions were 95°C for two-minutes, followed by 40 cycles of 95°C for 15 s and 60°C for 1 min. Relative expression levels were calculated using ΔΔCt method normalized to β-actin.

### Immunofluorescence Staining

Cardiac organoids were fixed in 4% paraformaldehyde (PFA) for 24-48 hours at 4°C, washed in PBS and processed for paraffin embedded sectioning. Organoids were embedded in paraffin blocks and sectioned at 5 μm thickness. For staining, sections were deparaffinized in Xylene and rehydrated through a graded ethanol series (100%-50%) and a final wash in distilled water (10 minutes per step). Antigen retrieval was performed in 10mM citric acid buffer (pH = 6.0) at 115 °C for 15-20 min, followed by cooling down at room temperature. Slides were then permeabilized in 0.1% Triton X-100 in PBS (PBS-T) and blocked in 5% normal goat serum in PBS-T and primary antibodies (Supplemental Table 1) were diluted in 1% normal-goat serum in PBS-T (0.1% triton) overnight at 4 °C in a humidified chamber. After three dPBS washes secondary antibodies were diluted in 1% NGS-PBST at room temperature in the dark for 1 hour. Nuclei were counterstained with DAPI 1 μg/μL (sigma). Slides were mounted with ProLong^TM^ Glass Antifade Mountant (Thermo Scientific) for imaging.

### Enzyme-Linked Immunosorbent Assay (ELISA) POSTN

Conditioned media were collected from cardiac organoid cultures at day 14 of maturation. Samples were centrifuged (10,000 x g for 5 min at 4°C) and stored at -80°C until experimentation. Secreted Periostin was quantified using the human POSTN ELISA kit (#ab213816, Abcam) following manufacturer’s instructions.

### Reactive Oxygen Species (ROS) in Organoid Lysates

Reactive oxygen species (ROS)/Reactive Nitrogen Species (RNS) were quantified using the DCF ROS/RNS assay kit (ab238535, Abcam) according to the manufacturer’s instructions. Organoid lysates were homogenized in PBS (10mg tissue mL^-1^), sonicated on ice and centrifuged at 10,000 x g for 5 min at 4 °C for debris removal. Fluorescence was measured at Ex/Em 480/530nm using a microplate reader (SpectraMax M3).

### Single-nuclei RNA Sequencing Analysis

Single-nuclei RNA isolation, library preparation and sequencing were performed by Novogene using the 10x Genomics Chromium Single Nuclei 3’ platform and Illumina NovaSEQ X. Single nuclei RNA sequencing (fastq) files were aligned to human transcriptomic reference (GRCh38), with CellRanger (v9.0.1) to generate filtered count matrices. Filtered count matrices were processed in Seurat (v5.2.1) for quality control, retaining nuclei with more than 500 unique molecular identifiers (UMIs), at least 200 detected features, and less than 5% mitochondrial gene content. Doublets were identified using DoubletFinder (v2.0.4), with the homotypic doublet proportion estimated and a presumed doublet rate of 8% per sample. Nuclei classified as doublets were excluded before proceeding with downstream integration analyses.

Sample integration was conducted with Harmony-based (v1.2.3) methodology implemented via Seurat utilizing 40 dimensions determined from the elbow plot following principal component analysis. Nearest neighbor graphs were computed and subsequently used for Louvain-based clustering identification with Seurat functions, *FindNeighbors* and *FindClusters,* respectively. UMAP dimensionality reduction was generated with the Seurat function *RunUMAP.* Cell Cycle Scoring was conducted with the Seurat function, CellCycleScoring, using reference “G2M” (n=54 genes) and “S” phase (n=43 genes).

Reference-based annotation of snRNA-seq integrated object was conducted using 10x single cell sequencing data from embryonic hearts (GSE106118)[52]. The single-cell fetal heart reference was down sampled to a maximum 2000 cells for each cell type. The reference then underwent *SCTransform*-based normalization with 3,000 variable features with Seurat. The single-nucleus cohort and reference were calculated using log normalization and 30 dimensions with the *FindTransferAnchors* function and subsequently used to label single nuclei with the *TransferData* function from Seurat. independently cross-validated against UCell-derived annotations using the Seurat functions *AddModuleScore* and *UCell*.

Following nuclei annotation, RNA raw read count matrices were converted into a *SingleCellExperiment* object (v1.28.1) for quality assessment using scuttle (v1.16.0). Outlier nuclei exceeding two median absolute deviations from the log_2-_transformed median count and genes detected in fewer than 10 nuclei were removed. Library size normalization and log-transformation were performed with *computeLibraryFactors* and *logNormCounts*, respectively. Normalized counts were aggregated by sample and cell type to generate pseudobulk matrices. Differential expression was performed using DESeq2 (v1.46.0) with Benjamini-Hochberg correction (α = 0.05) and log_2_ fold-change shrinkage via *lfcShrink* with the adaptive Student’s *t* prior (*apeglm*, v1.28.0). Variable genes were identified from *rawVars* (matrixStats, v1.5.0) on regularized log-transformed counts for principal component analysis using *plotPCA*. Data visualization was performed with ggplot2 (v3.5.1) and EnhancedVolcano (v1.24.0).

### Intracellular Calcium Assessment

Intracellular Ca^2+^ ([Ca^2+^]_i_) was measured using the Ca^2+^-sensitive dye Fura-2 and a dual-excitation (340/380 nm) spectrofluorometer (IonOptix LLC, Milton, MA, USA). First, cardiac organoids were incubated with 2.5 µmol/L Fura-2 for 20 minutes at room temperature, then washed with fresh regular Tyrode’s buffer containing (in mmol/L): 144 NaCl, 1 MgCl_2_, 10 HEPES, 5.6 glucose, 5 KCl, 1.2 NaH_2_PO4 (adjusted to a pH 7.4 with NaOH), and 1.5 CaCl_2_, for at least 10 minutes. Then, organoids were placed in a perfusion chamber adapted to the stage of an inverted Nikon eclipse TE2000-U fluorescence microscope. Fully intact organoids were kept steady in the perfusion chamber by a small, custom-built mesh. The organoids were superfused with Tyrode’s buffer at 37°C and Fura-2 fluorescence was acquired at an emission wavelength of 515±10 nm. The organoids were electric field-paced (20V) at different frequencies from 0.2 to 1.0 Hz.

The calibration was performed in cardiomyocytes “*ex vivo*” superfusing a free Ca^2+^ and then a Ca^2+^ saturating (5 mmol/L) solutions, both containing 10 µmol/L ionomycin (Sigma, St. Louis, MO) until reaching a minimal (R*_min_*) or a maximal (R*_max_*) ratio value, respectively. [Ca^2+^]_i_ was calculated as described previousl<u>y</u> using the following equation[53]:

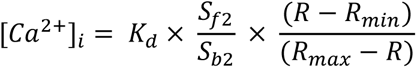

K*_d_* (dissociation constant) in was taken as 224 nmol/L. The scaling factors S*_f2_* and S*_b2_* were extracted from calibration as described Δ[Ca^2+^]_i_ [54] amplitude was defined as: peak [Ca^2+^]_i_ - resting [Ca^2+^]_i_.

### Spontaneous and Arrhythmic Activity Assessment

The incidence of autonomous Ca^2+^ firing activity was calculated for each group as the percentage of organoids showing spontaneous well-shaped Ca 2+ transients. The rate of autonomous activity was derived from the average peak-to-peak time (in seconds) as 1/t P-to-P, during a determined period. Pacing-induced arrhythmia was expressed as the percentage of organoids exhibiting any type of arrhythmic events based on intracellular Ca 2+ abnormalities, including early-after depolarization, delayed-after depolarization, tachyarrhythmia, etc. which do not correspond to an electric field stimulus during the pacing protocol. These were separated by the moment of predominant appearance in low-rate pacing (at 0.2 Hz) or during the entire protocol (0.2-1 Hz).

### Contractile dynamics analysis

Contractile activity of day 14 cardiac organoids was recorded by brightfield video microscopy under consistent imaging conditions. Videos were analyzed in Fiji (ImageJ) using the MuscleMotion plugin. Regions of interest encompassing beating tissue were selected, and motion traces were generated based on frame-to-frame pixel intensity changes. Contractile parameters, including amplitude and beat rate, were extracted and averaged per organoid for downstream analysis.

### Quantification of Rhythmic Motion Area

Brightfield videos were analyzed frame-by-frame using PIVlab v1.43 in MATLAB R2026b with custom MATLAB/Python processing. Following camera-motion correction and 15-pixel high-pass filtering, displacement was estimated by single-pass FFT cross-correlation. Sinusoidal regression identified rhythmic motion within 0.3-4 Hz, requiring at least three cycles. Rhythmic-motion area was expressed as a percentage of the entire segmented projected organoid, retaining unresolved tissue in the denominator. Assessable organoids without detected rhythm retained zero, technically immeasurable organoids were treated as missing.

### Drug Treatment

*LMNA*-mutant cardiac organoids were treated with nintedanib or pirfenidone to assess whether profibrotic signaling was therapeutically modifiable. Treatments were performed for a total of 7 days, with compounds added in two sequential treatment periods. The first dose was administered for 3 days, after which media was replaced, and a second dose was administered for an additional 4 days. Vehicle-treated organoids received 0.1% DMSO under the same media-change schedule. At the end of the 7-day treatment period, conditioned media was collected for POSTN ELISA, and organoids were harvested for western blot analysis or functional assessment.

### Human Fibrosis RT^2^ Profiler PCR Array

Fibrosis-associated gene expression was evaluated using the Human Fibrosis RT^2^ Profiler PCR Array (PAHS-120Z, QIAGEN), which profiles 84 genes associated with extracellular matrix remodeling, fibroblast activation, inflammatory signaling, and profibrotic pathways. Total RNA was isolated from *LMNA*-mutant, isogenic-corrected, and *LMNA*-mutant human induced pluripotent stem cell-derived cardiac organoids treated with 1 μM nintedanib for 7 days using the Quick-RNA Microprep Kit (R1050, Zymo Research). Two biological replicates (*n =2*) were analyzed per condition. Complementary DNA (cDNA) was synthesized using the iScript cDNA Synthesis Kit (1708891, Bio-Rad), and quantitative real-time PCR (qPCR) was performed using the RT^2^ Profiler PCR Array according to the manufacturer’s instructions. Cycle threshold (Ct) values were analyzed using the QIAGEN GeneGlobe RT^2^ Profiler PCR Array Data Analysis platform. Gene expression was normalized to ACTB, and relative expression was calculated using the comparative Ct (2^−ΔΔCt) method. Fold changes were determined by comparing *LMNA*-mutant organoids with isogenic-corrected controls and nintedanib-treated *LMNA*-mutant organoids with untreated *LMNA*-mutant controls. Expression profiles were examined to characterize fibrosis-associated transcriptional alterations and identify genes responsive to nintedanib treatment.

## FIGURE LEGENDS

**Supplemental Figure 1. Validation and quality control of *LMNA*-mutant and *LMNA*-corrected iPSC lines. (A)** RT-PCR analysis confirming clearance of Sendai virus (SeV) following reprogramming of *LMNA*-mutant iPSCs with *GAPDH* as the loading control. **(B)** Representative G-banded karyotypes demonstrating normal chromosomal integrity in *LMNA*-mutant and *LMNA*-corrected iPSC lines. **(C)** Representative brightfield images of control, *LMNA*-mutant, and *LMNA*-corrected iPSC-lines. **(D)** Immunofluorescence staining confirming pluripotency in control (SCT003-A), *LMNA*-mutant, and *LMNA*-corrected iPSCs for pluripotency markers *SOX2* and *SSEA-4* with DAPI marking nuclei. Merged images shown. **(E)** Directed differentiation assays of *LMNA*-mutant iPSCs by immunofluorescence staining of ectodermal markers (*PAX6, SOX1*), endodermal markers (*FOXA2, SOX17*), and mesodermal markers (*Brachyury, TBX6*). DAPI marks nuclei. Merged images shown. Scale bars = 100 µm.

**Supplemental Figure 2. Long-read RNA sequencing confirms absence of aberrantly spliced *LMNA* exon 6 length in control-iPSCs and human hearts. (A)** PacBio long-read RNA sequencing of control-iPSCs showing full-length *LMNA* transcript isoforms and corresponding TPM values. Canonical Lamin A and Lamin C isoforms are identified. **(B)** Quantification of the percentage of total *LMNA* TPM corresponding to transcripts with normal exon 6 length in publicly available PacBio long-read datasets from human control hearts (Pacbio Biosciences data repository).

**Supplemental Figure 3. Quantification of arrhythmic events. (A)** Autonomous calcium firing of organoids in Tyrode solution in beats per minute (bpm). **(B)** Percentage of organoids showing autonomous calcium firing in Tyrodes solution. **(C)** Percentage of organoids showing arrhythmic events in Tyrode solution and low-paced conditions (0.2-0.5 hz). **(D)** Percentage of organoids showing arrhythmic events in Tyrode solution.

**Supplemental Figure 4. Differentially Expressed Genes Across Individual Cardiac Lineages Identified by Single-nucleus RNA Sequencing.** Violin plots **(A-D)** depicting normalized expression levels of representative differentially expressed genes for the indicated cell populations in *LMNA*-mutant (blue) and isogenic *LMNA*-corrected (pink) organoids. Expression values are shown on a log-normalized scale. **(E)** Chord diagram depicting the top 20 unique endothelial cell-to-endothelial cell ligand-receptor interactions identified by CellChat analysis. **(F-J)** Dot plots indicating expression levels and % of cells expressing depicted genes, for the indicated cell types. **(K)** Quantification of contraction amplitude, **(L)** peak-to-peak interval, **(M)** time to peak, and **(N)** relaxation time in vehicle- and nintedanib-treated *LMNA*-mutant organoids.

## Notes

### Competing Interest Statement

The authors have declared no competing interest.

### Summary of Updates

This revised version includes additional Western blot analyses of total and phosphorylated p38 MAPK, quantification of rhythmic contractile activity following nintedanib treatment, and qPCR array profiling of fibrosis-associated genes. The manuscript and figures have been updated to incorporate these findings.

