## Supplementary figures and images for "*LMNA* Haploinsufficiency in Human iPSC-Derived Cardiac Organoids Reveals Early Fibrotic Signaling as a Therapeutically Targetable Process"

### Supplemental Figure 1

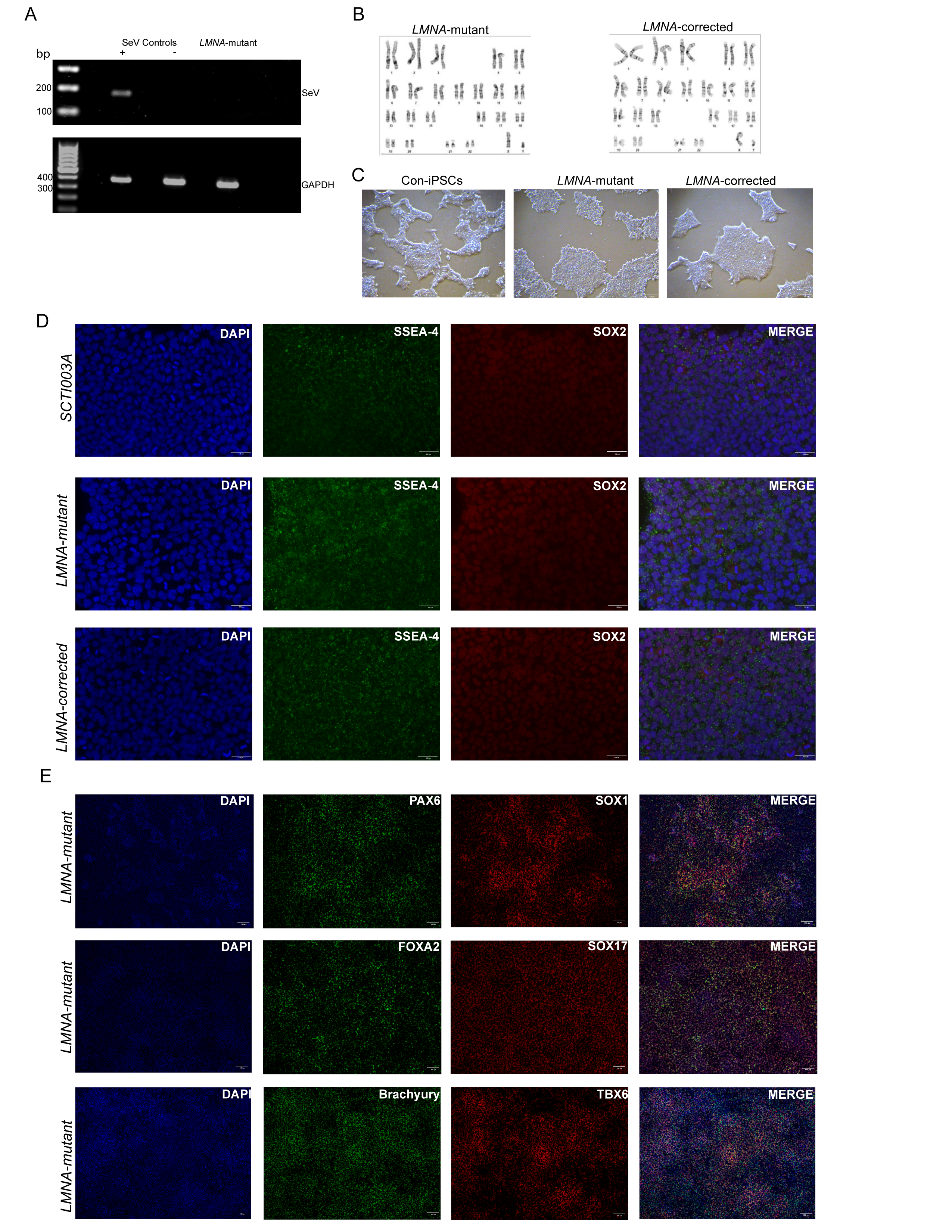

### Supplemental Figure 2

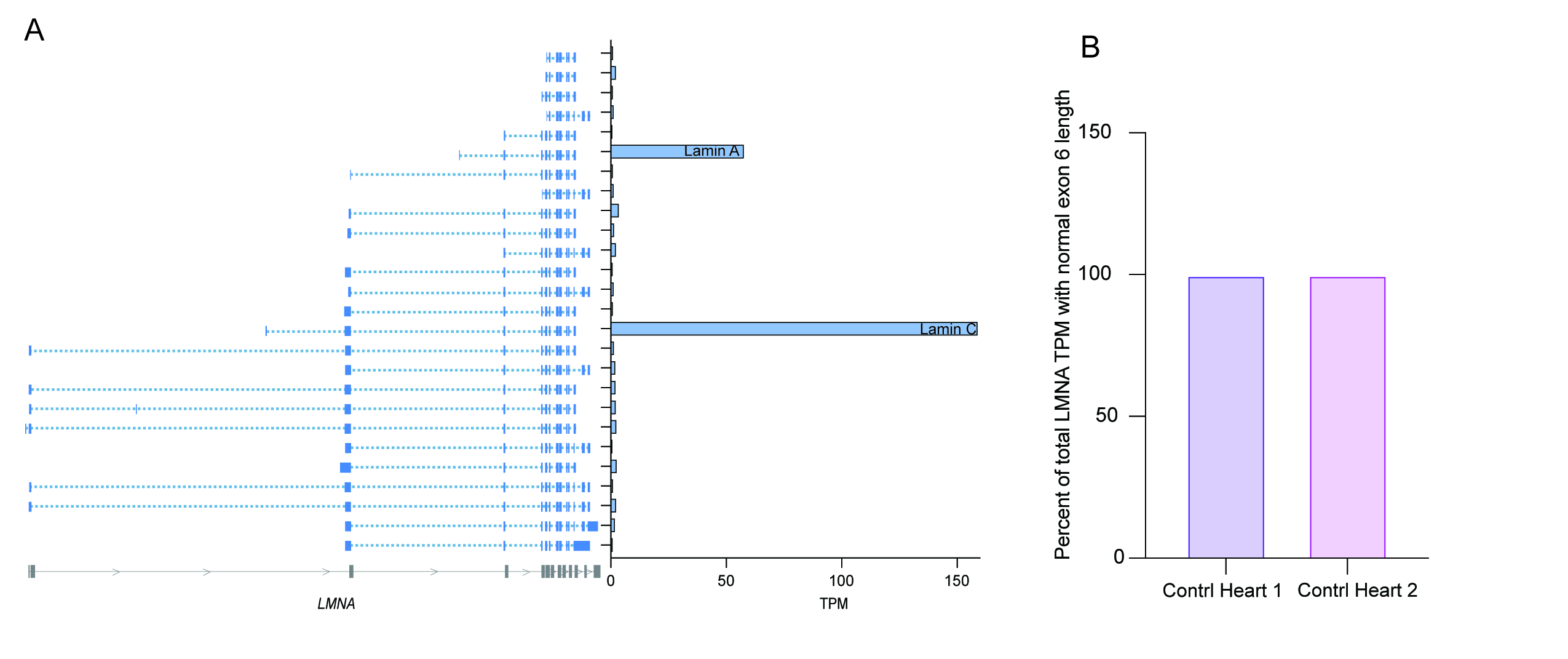

### Supplemental Figure 3

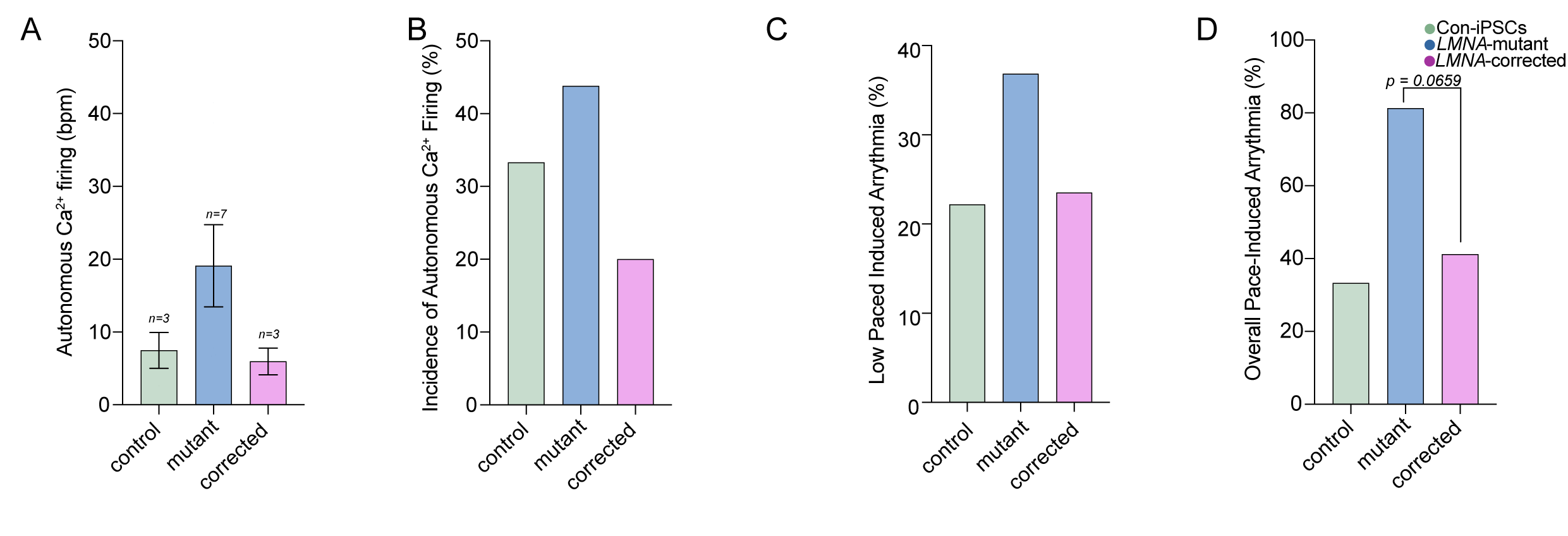

### Supplemental Figure 4

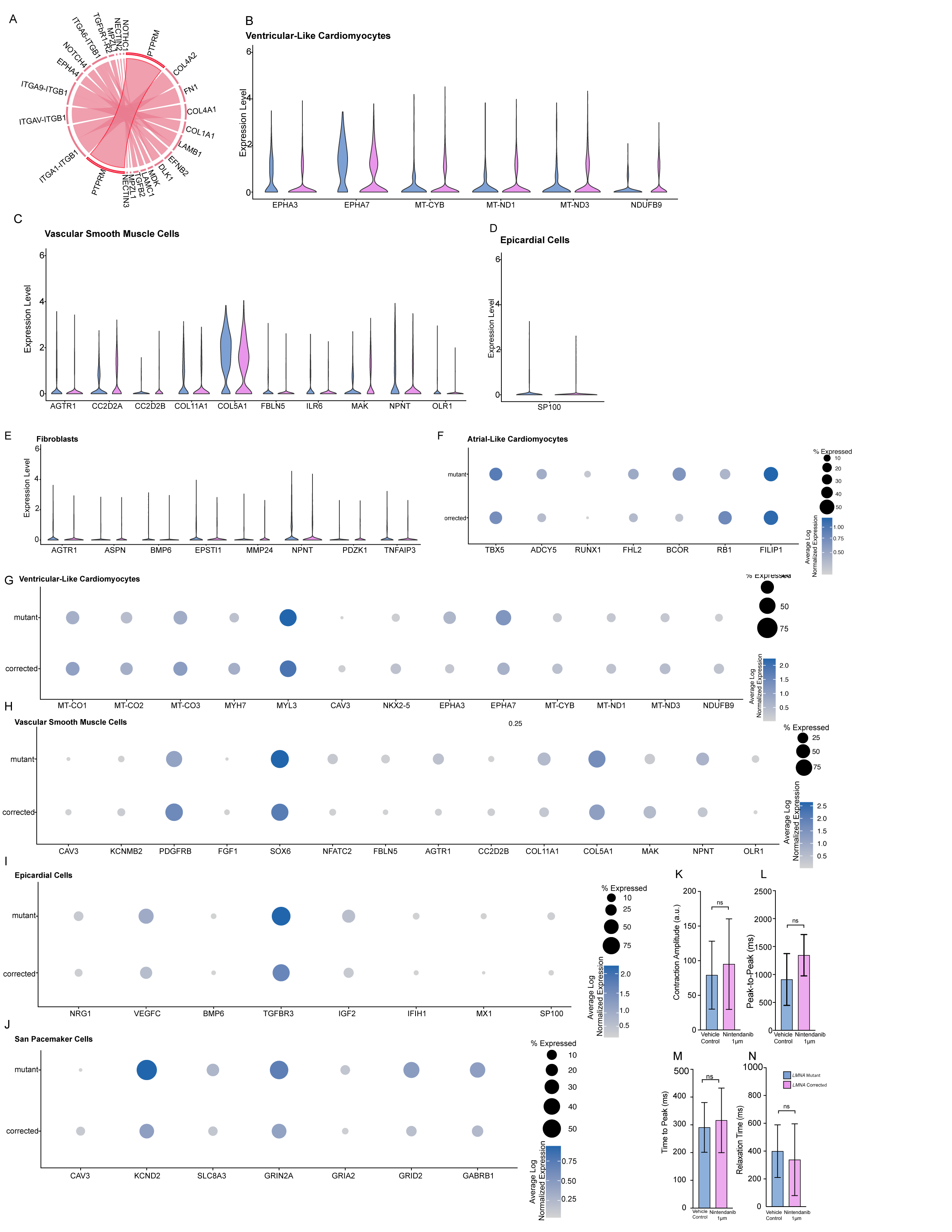
