## Supplemental Table 1 for "*LMNA* Haploinsufficiency in Human iPSC-Derived Cardiac Organoids Reveals Early Fibrotic Signaling as a Therapeutically Targetable Process"

| Antibody Name | Species | Vendor | Catalog Number | Dilution |
|---|---|---|---|---|
| P38 | Rabbit | Abcam | # 8690T | 1:500 |
| P-P38 | Rabbit | Abcam | # 4511T | 1:500 |
| POSTN | Mouse | R & D systems | #AF3548 | 1:500 |
| CTGF | Rabbit | Abcam | ab6992 | 1:500 |
| SOX-2 | Rabbit | Cell Signaling | #23064 | 1:400 |
| OCT-4 | Rabbit | Cell Signaling | #2750 | 1:200 |
| SSEA-4 | Mouse | Cell Signaling | #4755 | 1:400 |
| Lamin A/C (E-1) | Mouse | Santa-Cruz | SC-376248 | 1:500 |
| Cardiac Troponin | Mouse | Abcam | #AB8295 | 1 μg/mL |
| Beta Actin | Rabbit | Cell Signaling | #4970S | 1:1000 |
| Pax6 | Rabbit | Thermo Fisher | 42-6600 | 1:100 |
| Sox1 | Goat | R&D Systems | AF3369 | 1:50 |
| Brachyury | Rabbit | Abcam | ab20966 5 | 1:1000 |
| TBX-6 | Goat | R&D Systems | AF4744 | 1:100 |
| FOXA2 | Rabbit | Thermo Fisher | 701698 | 1:100 |
| SOX-17 | Goat | R&D Systems | AF1924 | 1:50 |
| Donkey IgG-Alexa Fluor Plus 488 | Anti-Rabbit | Thermo Fisher | A32790 | 1:1000 |
| Donkey IgG-Alexa Fluor Plus 647 | Anti-Goat | Thermo Fisher | A32849 | 1:1000 |
| Goat IgG-Alexa Fluor 594 | Anti-Rabbit | Invitrogen | #A11012 | 2 μg/mL |
| Goat IgG-Alexa Fluor 488 | Anti-Mouse | Invitrogen | #A11001 | 1 μg/mL |
