## Supplemental Table 2 for "*LMNA* Haploinsufficiency in Human iPSC-Derived Cardiac Organoids Reveals Early Fibrotic Signaling as a Therapeutically Targetable Process"

| Primer | Sequence |
| --- | --- |
| *LMNA* Off Target 2F | TGTAAAACGACGGCCAGTTGGAGCACTGAGTGATTAGC |
| *LMNA* Off Target 2R | CAGGAAACAGCTATGACCATCCCACATGTCCCTAATTCCC |
| *LMNA* Off Target 3F | TGTAAAACGACGGCCAGTACAAGGTGGACCGAATTTAGGA |
| *LMNA* Off Target 3R | CAGGAAACAGCTATGACCACATTAGCTCACTCACCCACA |
| *LMNA* Off Target 5F | TGTAAAACGACGGCCAGTCCATAAGTGCTCAATGCCAGC |
| *LMNA* Off Target 5R | CAGGAAACAGCTATGACCGGTGCCACCTCTGATAACCC |
