## Supplemental Table 3 for "*LMNA* Haploinsufficiency in Human iPSC-Derived Cardiac Organoids Reveals Early Fibrotic Signaling as a Therapeutically Targetable Process"

| Guide Sequence | PAM | Position | Strand | Donor Sequence (+strand; 5'->3') |
|---|---|---|---|---|
| CTTGGCTGCCAGTTGAAGGG | GGG | Chr1:156135890-156135912 | Minus | GTAGCCAGGTGTCTCCTACACCGACCCACGTCCCTCCTTCCCCATACTTAGGGCCCTTGGGAGCTCACCAAACCCTCCCACCCCCCTTCAGCTCGCTGCCAAGGAGGCGAAGCTTCGAGACCTGGAGGACTCACTGGCCCGTGAGCGGGACACCAGCCGGCGGCTGCTGGCGGAAAAGGAG |
